# Reply to *Cross-Contamination Explains ‘‘Inter and Intraspecific Horizontal Genetic Transfers’’ between Asexual Bdelloid Rotifers* (Wilson, Nowell & Barraclough 2018)

**DOI:** 10.1101/368209

**Authors:** Jean-François Flot, Nicolas Debortoli, Bernard Hallet, Jitendra Narayan, Karine Van Doninck

**Affiliations:** Evolutionary Biology & Ecology, Université libre de Bruxelles, 1050 Bruxelles, Belgium; Interuniversity Institute of Bioinformatics in Brussels – (IB)^2^, Bruxelles, Belgium; Laboratory of Evolutionary Genetics and Ecology, URBE, NAXYS, University of Namur, 5000 Namur, Belgium; Institut des Sciences de la Vie, Université Catholique de Louvain, 1348 Louvain-la-Neuve, Belgium

## Abstract

We thank Wilson et al. (2018) for their thorough re-analysis of our data and for their constructive criticisms that led our groups to exchange many stimulating emails over the last two years. Although we agree that inter-individual contamination can yield patterns suggestive of inter-individual recombination, we are not fully convinced by their criticisms of our 2016 dataset and would like to point here briefly to some inaccuracies and likely errors in their interpretation of our chromatograms.

---

We thank Wilson *et al*. (2018) for their thorough re-analysis of our data and for their constructive criticisms that led our groups to exchange many stimulating emails over the last two years. Although we agree that inter-individual contamination can yield patterns suggestive of inter-individual recombination, we are not fully convinced by their criticisms of our 2016 dataset and would like to point here briefly to some inaccuracies and likely errors in their interpretation of our chromatograms (available at https://github.com/jflot/Debortoli2016CurrentBiology).

Wilson *et al*.’s criticism of our results rests on two main arguments: based on their ConTAMPR analysis of the pattern of minor peaks in some of our chromatograms (often barely distinguishable from background noise), they interpret our proposed inter-specific recombination patterns as merely the result of cross-contamination, namely the inadvertent co-extraction of several individuals in the same tube; and based on their finding of triple peaks in some other chromatograms they conclude that our proposed intra-specific recombination patterns are also caused by cross-individual contamination.

If we inadvertently co-extracted several individuals at once as suggested by Wilson *et al*., we should observe superposition of several sequences of approximately equal intensities in at least some of our COI chromatograms (the sequencing of which did not involve any whole-genome amplification step), especially given that rotifers are eutelic (hence juveniles and adults contain strictly the same amount of DNA). This was never the case: the minor peaks were always much smaller than the main ones. This appears more compatible with the hypothesis of minute amounts of carry-over or post-PCR contamination than with co-extraction of several individuals. It is well known that there is tiny amounts of DNA ‘floating around’ in any laboratory (Gruber *et al*. 2015); background-level contamination may also occur post-PCR, during cycle-sequencing and capillary electrophoresis. Alternatively, contamination by the gut content of the rotifers we sequenced (which might contain traces of ingested DNA from dead rotifers) cannot be ruled out and could explain some of the minor peaks observed. These possible explanations for the minor peaks incriminated by Wilson *et al*. were not considered in their article.

To see whether the minor peaks in the COI chromatograms of our 2016 article were unusually abundant, we compared them with those of another published COI dataset of groundwater amphipods (Flot *et al*. 2012). These amphipods are several millimetres long so the chances of putting by mistake two individuals in the same tube are virtually nil. As we found the ConTAMPR approach of Wilson *et al*. error prone and time consuming when performed manually, we implemented it in an automatic program dubbed autoConTAMPR (https://github.com/jnarayan81/autoConTAMPR; Narayan *et al*., in prep.). When using autoConTAMPR to test for contamination in each COI chromatogram pair from our Current Biology dataset (instead of focusing solely on individuals for which we detected patterns of inter-specific transfers, as Wilson *et al*. did using their manual approach), we found statistically significant hits (P<1%) for 49 of the 80 individuals for which both chromatograms were available (for two of them, the chromatogram obtained using primer HCO2198 was of low quality and had to be discarded), i.e. 61%; for the amphipod dataset, it was 42/67 individuals, i.e. 63%. This indicates that the level of background contamination detected by autoConTAMPR in our Current Biology dataset is not unusual for COI chromatograms and can be parsimoniously explained by small amounts of carry-over or post-PCR contamination (instead of several individuals being inadvertently extracted in the same tube).

We also analysed separately the forward and reverse chromatograms (instead of combining information from both as in Wilson *et al*.). For our Current Biology dataset we found 41 P<1% hits (51%) in the chromatograms obtained using primer LCO1490 and 18 P<1% hits (22.5%) in those obtained using primer HCO2198, whereas only four individuals (B11, A3B1, Hprim12 and H4–04; 5%) had P<1% hits on the same sequence for both chromatograms; for the amphipod dataset the figures were 67% for LCO1490, 25% for HCO2198 and 6% with at least one hit identical on both. Again, these percentages were strikingly similar between the two COI datasets, and the rotifer sequences appeared actually less frequently contaminated than the amphipod ones… Only one of the six rotifers for which we inferred putative interspecific gene transfers (B11) had identical P<1% hits on both chromatograms: these were matches with species D and E, whereas the inferred horizontal transfer was from E to A. Regarding the other five individuals: B39 (inferred mitochondrial capture from C to E) had no P<1% hit; B22 (inferred transfers from E to C) had a P<1% hit with species F for primer LCO1490 but none with species E; B3B1 (inferred transfer from E to A) had P<1% matches with species E and D for LCO1490; B14 (inferred transfers from C and E to A) had P<1% matches with species D and E for HCO2198 but none with species C; and D14 (inferred transfer from C to A) had P<1% matches with species B, D, E and F for LCO1490 but none with species C. Hence, although we cannot rule out at this stage that some of the signatures of interspecific recombination reported in our 2016 article actually resulted from contamination, the evidence at hand is far less conclusive than asserted by Wilson *et al*. and certainly not sufficient to discard our results.

Concerning intraspecific recombination, on the other hand, Wilson *et al*.’s assertion that the patterns we observed result from contamination clearly does not hold. Here Wilson et al. did not use ConTAMPR but instead observed a few triple peaks in the vicinity of single-nucleotide polymorphism (SNP) positions in our EPIC63 chromatograms and wrote: ‘No matter how they are shifted, though, two alleles cannot yield triple peaks. To find triple peaks therefore demonstrates the presence of three or more alleles, and thus DNA from more than two individuals’. This assertion is not true: in the case of length-variant heterozygotes (Flot *et al*. 2006), PCR-induced recombination among alleles (Judo *et al*. 1998; Cronn *et al*. 2002) does yield triple peaks when an incompletely elongated PCR product re-anneals with a fragment originating from the other haplotype during subsequent PCR cycles, creating chimeric haplotypes (Fig. 1).

**Fig. 1.**
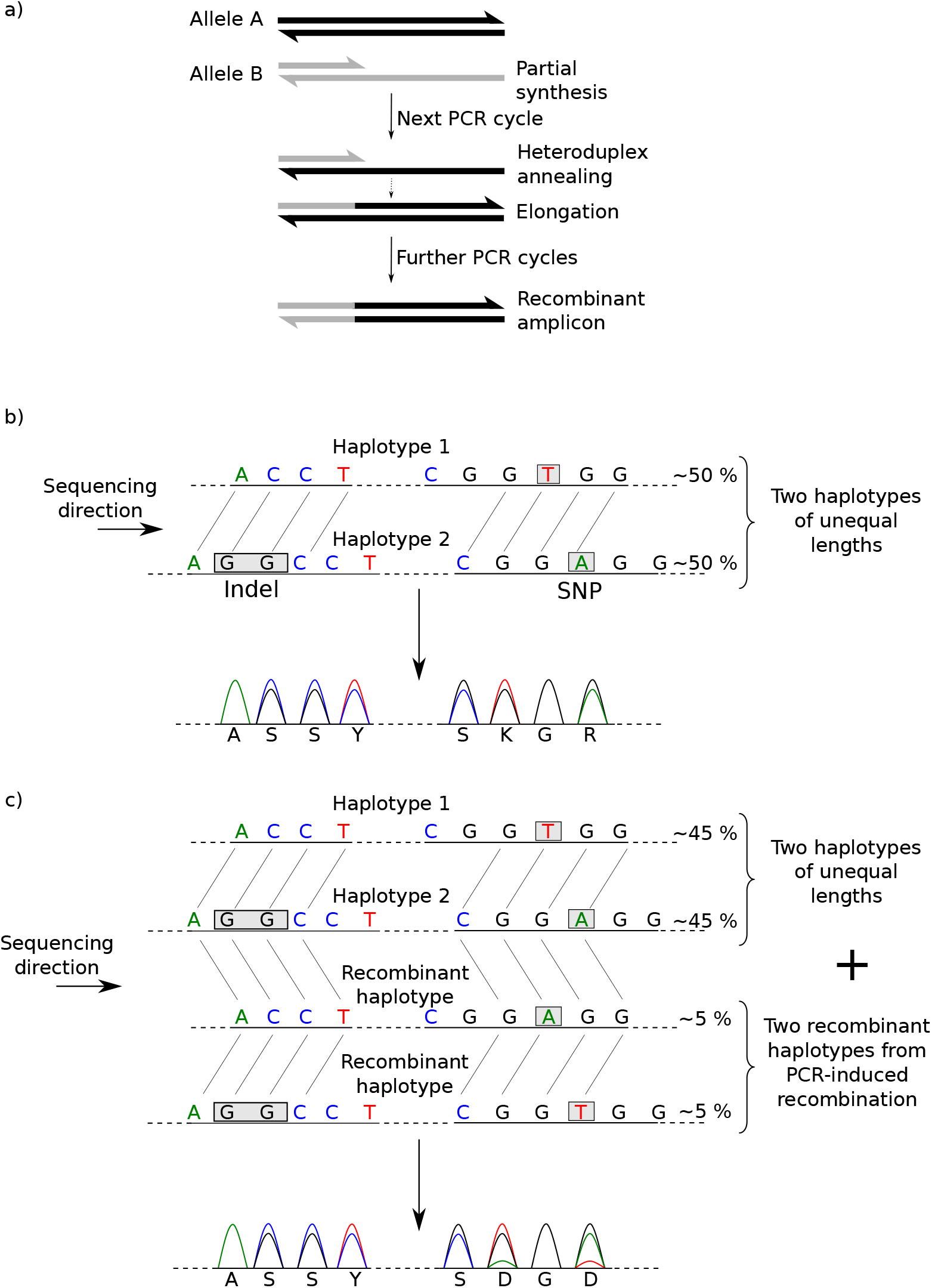
PCR-induced recombination in length-variant individuals yields triple peaks. a) When incomplete PCR products anneal with a fragment originating from the other haplotypes and are elongated by the *Taq* polymerase, recombinant amplicons are produced, resulting in chimeric haplotypes. b) In the absence of PCR-induced recombination, the chromatograms of length-variant heterozygotes comprise only single and double peaks, but no triple peaks. c) In the presence of recombinant haplotypes resulting from PCR-induced recombination, triple peaks are observed.

Based on Fig. 1, several predictions can be made regarding the triple peaks generated by PCR-induced recombination in length-variant heterozygotes: 1) they are only found downstream of the heterozygous indel, in the vicinity of a SNP visible on the other strand; 2) there are at most two triple peaks for each SNP visible on the other strand; 3) the distance between the two triple peaks produced by a given SNP is equal to the length difference of the two ‘real’ haplotypes; and 4) as the probability of PCR-induced recombination occurring between two loci increases with the distance separating then, the third peaks for SNPs close to the indel are barely visible but those farther away from the indel are more conspicuous. Careful scrutiny of our chromatograms verified these four predictions in all three individuals incriminated by Wilson *et al*. (H4–28, D22 and D23), highlighting that PCR-induced recombination is a more parsimonious explanation for these triple peaks than the presence of contaminant DNA. We also confirmed this hypothesis by cloning and sequencing some of our PCR products, which revealed the presence of recombinant haplotypes (data not shown).

We are presently replicating our study without whole-genome amplification step, by sequencing clonal lineages obtained from each sampled individual. Our preliminary results from this study (Debortoli *et al*., in prep.) confirm the occurrence of trios of allele-sharing individuals, suggesting that intra-specific transfers do occur among bdelloids. Another project using RADseq is currently under progress and will specifically look for evidence of inter-specific transfers. In the meantime, the question whether or not bdelloids exchange genes in unusual ways (as proposed independently by Signorovitch *et al*. 2015 and by Debortoli *et al*. 2016) remains open and will certainly lead to exciting new discoveries.

